# Memory B cells control SARS-CoV-2 variants upon mRNA vaccination of naive and COVID-19 recovered individuals

**DOI:** 10.1101/2021.06.17.448459

**Authors:** Aurélien Sokal, Giovanna Barba-Spaeth, Ignacio Fernández, Matteo Broketa, Imane Azzaoui, Andrea de La Selle, Alexis Vandenberghe, Slim Fourati, Anais Roeser, Annalisa Meola, Magali Bouvier-Alias, Etienne Crickx, Laetitia Languille, Marc Michel, Bertrand Godeau, Sébastien Gallien, Giovanna Melica, Yann Nguyen, Virginie Zarrouk, Florence Canoui-Poitrine, France Noizat-Pirenne, Jérôme Megret, Jean-Michel Pawlotsky, Simon Fillatreau, Pierre Bruhns, Felix A. Rey, Jean-Claude Weill, Claude-Agnès Reynaud, Pascal Chappert, Matthieu Mahévas

**Affiliations:** Institut Necker Enfants Malades (INEM), INSERM U1151/CNRS UMS 8253, Université de Paris, Paris, France; Service de Médecine Interne, Centre Hospitalier Universitaire Henri-Mondor, Assistance Publique-Hôpitaux de Paris (AP-HP), Université Paris-Est Créteil (UPEC), Créteil, France; Institut Pasteur, Unité de Virologie Structurale, CNRS UMR 3569, Paris, France; Institut Pasteur, Unité Anticorps en thérapie et pathologie, UMR 1222 INSERM, France; INSERM U955, équipe 2. Institut Mondor de Recherche Biomédicale (IMRB), Université Paris-Est Créteil (UPEC), Créteil, France; Département de Virologie, Bactériologie, Hygiène et Mycologie-Parasitologie, Centre Hospitalier Universitaire Henri-Mondor, Assistance Publique-Hôpitaux de Paris (AP-HP), Créteil, France; INSERM U955, équipe 18. Institut Mondor de Recherche Biomédicale (IMRB), Université Paris-Est Créteil (UPEC), Créteil, France; Service de Maladies Infectieuses, Centre Hospitalier Universitaire Henri-Mondor, Assistance Publique-Hôpitaux de Paris (AP-HP), Université Paris-Est Créteil (UPEC), Créteil, France; Service de Médecine interne, Hôpital Beaujon, Assistance Publique-Hôpitaux de Paris, Université de Paris, Clichy, France; Département de Santé Publique, Unité de Recherche Clinique (URC), CEpiA (Clinical Epidemiology and Ageing), EA 7376- Institut Mondor de Recherche Biomédicale (IMRB), Centre Hospitalier Universitaire Henri-Mondor, Assistance Publique-Hôpitaux de Paris (AP-HP), Université Paris-Est Créteil (UPEC), Créteil, France; Etablissement Français du Sang (EFS) Ile de France, Créteil, France; Plateforme de Cytométrie en Flux, Structure Fédérative de Recherche Necker, INSERM US24-CNRS UMS3633, Paris, France; Inovarion, Paris, France

## Abstract

How a previous SARS-CoV-2 infection may amplify and model the memory B cell (MBC) response elicited by mRNA vaccines was addressed by a comparative longitudinal study of two cohorts, naive individuals and disease-recovered patients, up to 2 months after vaccination. The quality of the memory response was assessed by analysis of the VDJ repertoire, affinity and neutralization against variants of concerns (VOC), using unbiased cultures of 2452 MBCs. Upon boost, the MBC pool of recovered patients selectively expanded, further matured and harbored potent neutralizers against VOC. Maturation of the MBC response in naive individuals was much less pronounced. Nevertheless, and as opposed to their weaker neutralizing serum response, half of their RBD-specific MBCs displayed high affinity towards multiple VOC and one-third retained neutralizing potency against B.1.351. Thus, repeated vaccine challenges could reduce these differences by recall of affinity-matured MBCs and allow naive vaccinees to cope efficiently with VOC.

## Introduction

The COVID-19 pandemic caused by severe acute respiratory syndrome coronavirus 2 (SARS-CoV-2) has resulted in more than 165 million infections and at least 3.5 million deaths. Vaccination represents the main hope to control the pandemic. COVID-19 vaccines containing nucleoside-modified mRNA encoding the original Wuhan-Hu-1 SARS-CoV-2 spike glycoprotein (S) developed by Pfizer/BioNTech (BNT162b2) and Moderna (mRNA-1273) are now being deployed worldwide. They were shown to be safe, highly effective to prevent infection and to control disease severity (Baden et al., 2021; Dagan et al., 2021; Polack et al., 2020).

The emergence of SARS-CoV-2 variants bearing mutations in key B cell epitopes, however, has raised concerns that viral evolution will erode natural immunity or the protection offered by vaccination. One early mutation in the spike protein (D614G), which shifts the equilibrium between the open and closed protein conformation without modifying antibody neutralization, has become globally dominant (Plante et al., 2021; Weissman et al., 2021; Yurkovetskiy et al., 2020). Since then, novel variants of concern (VOC) or of interest (VOI) have spread around the world, with additional combinations of mutations and deletions mainly located in the ACE-2 receptor-binding domain (RBD) and the N-terminal domain of the S protein. Mutations in the RBD are of particular importance, since a large fraction of neutralizing antibodies elicited after infection and vaccination target this domain. The selective advantage provided by these mutations has resulted in their increasing prevalence: N501Y in the B.1.1.7 (alpha) variant; K417N, E484K, N501Y in the B.1.351 (beta) variant, K417T, E484K, N501Y in the P.1 (gamma) variant and L452R, E484Q, or L452R, T478K in the B1.617.1 (kappa) and B1.617.2 (delta) variants, respectively (Cherian et al., 2021; Davies et al., 2021; Greaney et al., 2021a; Tegally et al., 2021).

Higher infectiousness of the B.1.1.7 variant does not impair neutralizing antibody response (Davies et al., 2021; Garcia-Beltran et al., 2021; Planas et al., 2021; Supasa et al., 2021). By contrast, E484K and K417T/N mutations in the B.1351 and P.1 strains markedly reduced the neutralization potency in COVID-19 recovered or naive vaccinated individuals (Cele et al., 2021; Edara et al., 2021; Greaney et al., 2021a; Hoffmann et al., 2021; Planas et al., 2021; Wang et al., 2021a; Xie et al., 2021). Even though infection with VOC/VOI remain possible after successful vaccination (Hacisuleyman et al., 2021), the effectiveness of the BNT162b2 vaccine against B.1.351 in preventing severe disease was recently demonstrated during the vaccination campaign in Qatar (Abu-Raddad et al., 2021).

In addition to the serum IgG response, the generation of memory B cell (MBC) against the SARS-CoV-2 virus represents another layer of immune protection (Dugan et al., 2021; Gaebler et al., 2021; Rodda et al., 2021; Sakharkar et al., 2021; Sokal et al., 2021). MBCs not only persist after infection but continuously evolve and mature by progressive acquisition of somatic mutations in their variable region genes to improve affinity through an ongoing germinal center response, potentially driven by antigenic persistence (Gaebler et al., 2021, 2021; Rodda et al., 2021; Sokal et al., 2021). MBCs further drive the recall response after antigenic rechallenge by differentiating into new antibody-secreting cells (ASC) displaying the diverse array of high-affinity-antibodies contained in the MBC repertoire.

The remarkable convergence of the anti-RBD response across COVID-19 recovered and naive vaccinated individuals shaped by recurrent germline gene families may favor viral mutational escape as one single mutation in the RBD can confer a selective advantage by reducing binding and neutralizing activity of antibodies (Garcia-Beltran et al., 2021; Greaney et al., 2021a). This can be theoretically counter-balanced by the diversity of the MBC repertoire, which contains a broad variety of clones with different specificities. Upon vaccination, COVID-19 recovered patients showed a striking expansion of RBD-specific MBCs (Goel et al., 2021; Wang et al., 2021b) and elicited a strong serum antibody response including cross-neutralizing antibodies against VOC (Ebinger et al., 2021; Konstantinidis et al., 2021; Manisty et al., 2021; Saadat et al., 2021; Samanovic et al., 2021; Stamatatos et al., 2021; Wang et al., 2021b). Much less is known about the long-term stability, dynamics and functionality of the MBC repertoire after repeated antigenic stimulation. How the MBC pool will contract or expand its diversity after a new challenge is of major importance in the context of vaccination schemes with repeated homologous or heterologous booster doses and coexistence of multiple variants of concerns. Here, we longitudinally characterized the dynamics, clonal evolution, affinity and neutralization capacity of anti-SARS-CoV-2 memory B cell response after mRNA vaccination in naive and SARS-CoV-2 recovered individuals from the analysis of over two thousand naturally expressed antibodies from single-cell cultured RBD-specific MBCs.

Our study explores the intrinsic diversity of the MBC repertoire selected against RBD protein upon infection and/or vaccination and its capacity to cope with multiple variants of SARS-CoV-2. We demonstrate that mRNA vaccination selects high-affinity neutralizing clones without compromising the overall MBC pool.

## Results

### mRNA vaccination boosts serum IgG levels in SARS-CoV-2-recovered and naive individuals

The B cell immune response elicited by vaccination was compared between two cohorts, one of patients previously infected with SARS-CoV-2 six months to one year ago and one of virus-naive individuals having received, respectively, one or two doses of the Pfizer-BioNTech mRNA vaccine (BNT162b2). We previously characterized the longitudinal evolution and maturation, up to 6 months after infection, of SARS-CoV-2-responding B cells in a cohort of mild ambulatory (M-CoV) and severe forms of COVID-19 requiring oxygen (S-CoV) (Sokal et al., 2021). These patients have all been infected during the first pandemic wave in France. Thirty-four patients from this cohort, further referred to as SARS-CoV-2 recovered, were sampled at 12 months (**see Table S1)**. At that time, 8 patients had already received one dose of the BNT162b2 vaccine, 26 had not yet been vaccinated. Among these 26, 11 received one dose after the 12-month visit. This initial cohort was further extended by the enrollment of 9 additional mild COVID-19 patients, which received one dose of the vaccine. For the second cohort, we recruited 25 healthcare workers with no clinical history of COVID-19 and no serological evidence of previous SARS-CoV-2 infection (naive individuals) receiving 2 doses of vaccines as part of the French vaccination campaign.

For SARS-CoV2 recovered patients, the vaccine dose was administered in median 309 days post COVID-19 symptom onset (range: 183-362 days, **see Table S1**) and was referred to as vaccine “boost”. For SARS-CoV-2 naïve patient the second vaccine dose, also referred to as “boost”, was received in median 28 days after initial priming. Post-boost blood samples were collected from all vaccinated participants in median 8 days (range 5-22 days) and 2 months after the injection (**Figure 1A** and **see Table S1**). Eight SARS-CoV-2 naive donors were additionally sampled after the first dose of vaccine, referred to as “prime”.

**Figure 1.**
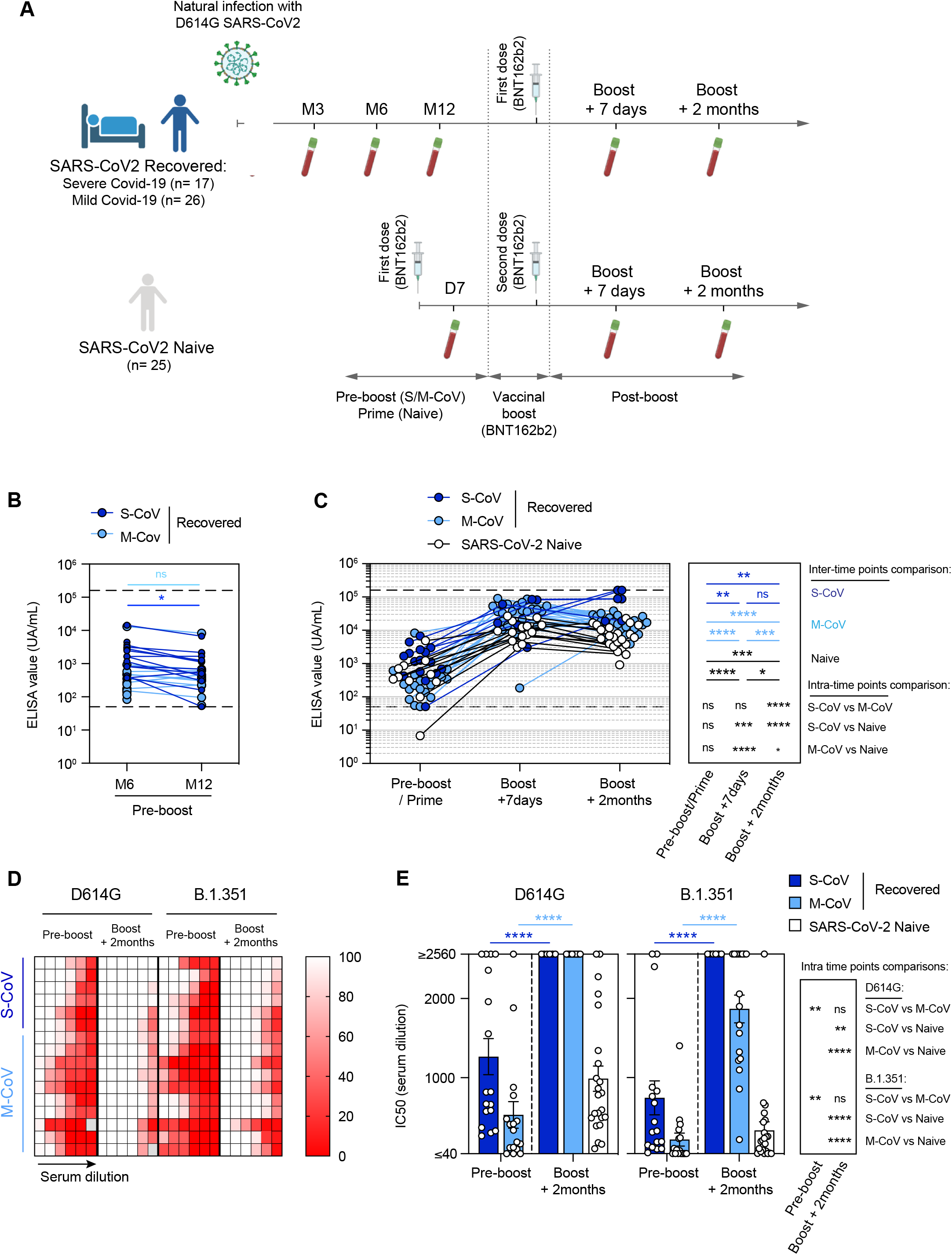
Longitudinal characterization of the humoral response against SARS-CoV-2 in SARS-CoV-2 recovered and naive patients after BNT162b2 vaccination. **(A)** Cohort design **(B)** Anti-SARS-CoV-2 RBD serum IgG level measured by ELISA in 31 SARS-CoV-2 recovered patients (S-CoV; dark blue; M-CoV: light blue) at 6 (M6) and 12 months (M12) post symptom onset. **(C)** Evolution of the anti-SARS-CoV-2 RBD serum IgG level after BNT162b2 vaccination. ELISA values are shown at pre-boost (M6 or M12) for SARS-CoV-2 recovered (S-CoV: dark blue; M-CoV: light blue) or after the first dose (Prime) for naive patients (white), as well as 7 days and 2 months after the vaccine boost. The lower dashed line indicates the positivity threshold and the upper dashed line indicates the upper limit of detection as provided by the manufacturer **(D)** Heatmap representing the observed in vitro neutralization of SARS-CoV-2 D614G (left) and B.1.351 variant (right) viruses by sera from SARS-CoV-2 recovered patients at the pre-boost and boost + 2 months time points (serial dilutions: 1/10, 1/40, 1/160, 1/640, 1/2560, 1/10240). Each line represents one patient. **(E)** Half maximal inhibitory concentration (IC50) for all sera tested from SARS-CoV-2 recovered and naive patients at pre-boost and boost + 2 months time points against SARS-CoV-2 D614G and B.1.351 viruses. Bars indicate mean±SEM. (B) Two-way ANOVA with multiple comparisons of all groups; (C) Repeated measures mixed effects model analysis with two sets of multiple comparisons (between donor groups inside each time-point and between time points for each donor group) and (E) repeated measures mixed effects model analysis with multiple comparisons between time points for each Recovered donor group and Kruskal-Wallis with multiple comparisons between donor groups inside each time-point were performed (Benjamini, Krieger and Yekutieli FDR correction was used for all multiple comparisons). (****P<0.0001, *** P<0.001, **P < 0.01, *P < 0.05). See also Figure S1 and Table S1.

We first measured the pre- and post-boost evolution of WT RBD IgG serum titers in SARS-CoV-2 recovered patients. Anti-RBD IgG levels remained remarkably stable between 6 (M6) and 12 months (M12) post-infection, with only a mild decrease seen in S-CoV patients (**Figure 1B**). In line with previous reports, a single dose of mRNA vaccine elicited a strong recall response in all patients, with anti-RBD IgG titers increasing on average by 24-fold in S-CoV and 53-fold in M-CoV patients as compared with pre-boost titers (**Figure 1C**).

In SARS-CoV-2 naive individuals, the mRNA vaccine boost also induced a robust anti-RBD IgG response (average of 25-fold-increase) although titers remained inferior to SARS-CoV-2-recovered patients at all time points (mean 10,870 vs. 75,511 UA/mL in S-CoV and 55,024 in M-CoV, *P*-value 0.005 and <0.0001, respectively). Despite contraction of the response in all groups, anti-RBD IgG were at least 10-fold higher than pre-boost/prime levels at 2 months post boost (**Figure 1C**).

To further assess the neutralizing activity against circulating variants of concern (VOC) in sera from SARS-CoV-2 recovered and naive individuals, we performed neutralization assays, using a focus reduction neutralization test with authentic SARS-CoV-2 viruses carrying the dominant Spike protein D614G amino acid change in the S1 domain of the S protein, and on the B.1.351 VOC harboring three mutations in the RBD (N501Y, which increases the affinity for the ACE2 receptor, E484K and K417N, which are implicated in escape from neutralizing antibodies (Harvey et al., 2021)), along with several mutations in other Spike domains. Twelve months after infection, all sera from recovered patients demonstrated neutralization potential against the D614G SARS-CoV-2 strain, which was more pronounced in S-CoV than in M-CoV (**Figure 1D-E and Figure S1**). As expected, this potential was similarly reduced in both groups against the B.1.351 VOC strain. As previously reported (Goel et al., 2021; Reynolds et al., 2021; Stamatatos et al., 2021; Wang et al., 2021b), a boost mRNA vaccine strongly enhanced the overall neutralizing potency of SARS-CoV-2 recovered patients against both the D614G and the B.1.351 viruses, with all S-CoV and M-CoV sera achieving IC50 >1/2560 for the D614G strain. All sera from S-CoV patients reached a similar neutralization level against the VOC B.1.351 strain. For M-CoV patients, the gain in neutralization potency against B.1.351 was also clear albeit more modest (>3 fold increased IC50 compared to pre-boost levels). Sera from all naive individuals also showed detectable neutralizing activity against the D614G strain at the final time point in our study (mean IC50: 1/988, range ≥2560 to 1/103), although never reaching the levels seen in SARS-CoV-2 recovered patients. This potency was further reduced against the VOC B.1.351 strain (mean IC50 1/332), with 8 out of 23 patients (35%) displaying IC50 inferior below 1/100 against this variant.

Together these data show that a boost of mRNA vaccine based on the Wuhan-Hu-1 S protein induces a strong recall response in all SARS-CoV-2 recovered patients, with strong neutralizing IgG serum levels against both D614G and B.1.351 VOC in most individuals. Despite a similar magnitude of the boost IgG response, the neutralization potency of the antibodies generated, however, appeared lower in naive individuals than in SARS-CoV-2 recovered patients, suggesting a key role for the matured memory B cell pool present in SARS-CoV2 recovered patients in modeling the quality of the vaccine response, including the cross-neutralization of VOC.

### mRNA vaccination mobilizes the memory B cell pool in SARS-CoV-2-recovered patientss

To track the dynamics of the overall SARS-CoV-2-specific B cell response after mRNA vaccine boost, we used successive staining with His-tagged RBD from the Wuhan-Hu-1 strain and two fluorescently labeled anti-His antibodies to stain SARS-CoV-2 RBD-specific B cells (**Figure 2A and Figure S2A**). We focused on RBD recognition because it represents a large fraction of the virus neutralization potency, but we also analyzed S-specific cells in all samples, using a similar staining strategy. The percentage of RBD-specific and S-specific CD27^+^IgD^-^ B cells appeared remarkably stable between 6 and 12 months in SARS-CoV-2 recovered patients, irrespectively of initial disease severity, thus confirming the generation of germinal-center-derived long-lived memory B cell after natural infection **(Figure 2B and Figure S2B)**. RBD-specific memory B cells (MBCs) significantly expanded after one dose of mRNA vaccine before a modest contraction at 2 months (**Figure 2C and Figure S2C**). In contrast, only low numbers of RBD-specific B cells were detectable in naive individuals after the first dose with a non-significant impact of boost vaccination (**Figure 2C**), although RBD-specific ASCs were observed in all donors in the early steps of the post-boost response **(Figure S2D)**. The frequency of RBD-specific MBCs persisting 2 months after the boost in naive individuals remained significantly lower than the levels of MBCs observed in SARS-CoV-2 recovered patients before vaccination (**Figure 2C**), with similar profiles observed for spike-specific memory B cells **(Figure S2E)**.

**Figure 2.**
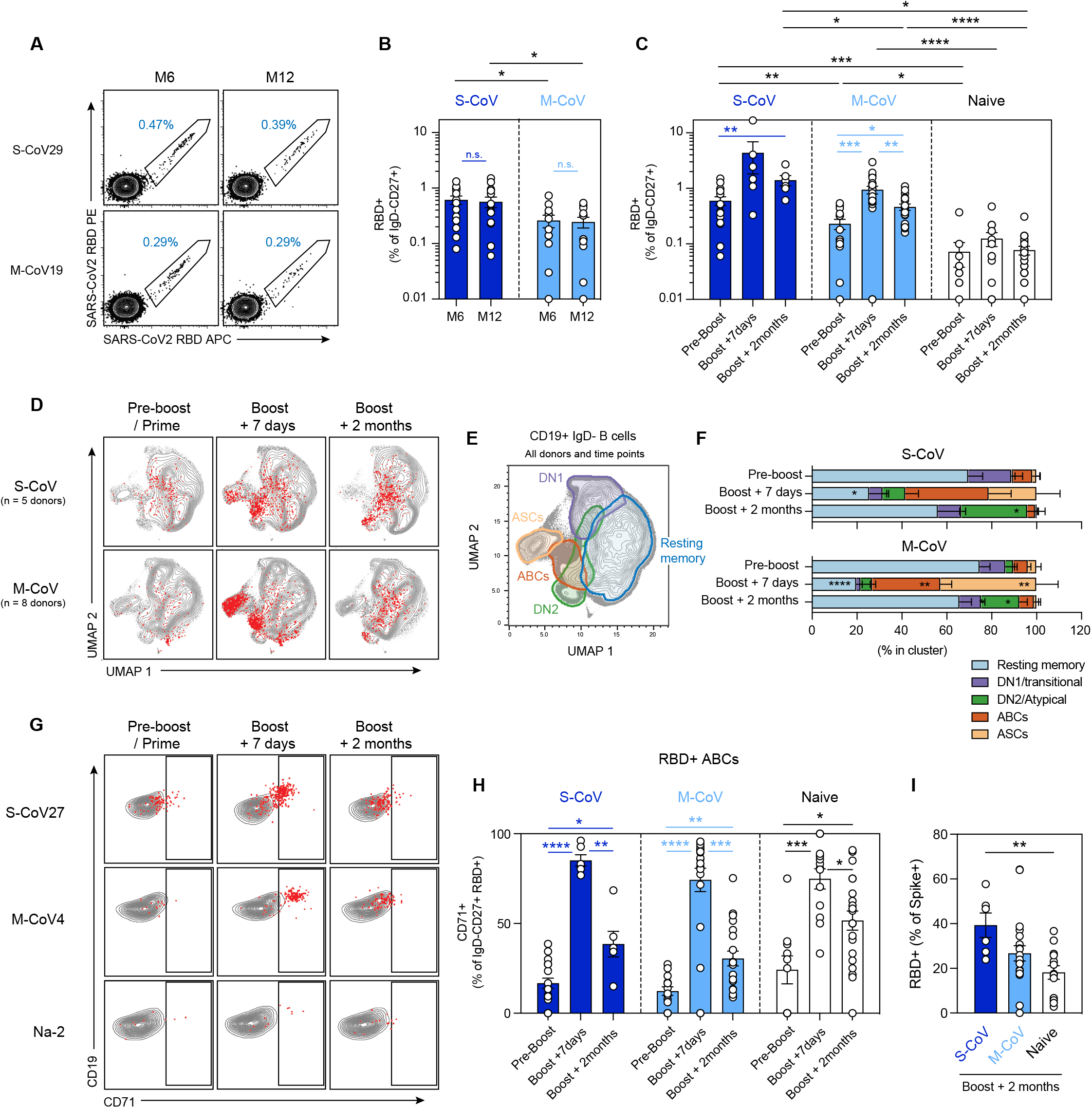
Phenotypic evolution of SARS-Cov-2 RBD-specific B cells after vaccination in SARS-CoV-2 recovered and naive patients. **(A)** Representative dot-plot of SARS-CoV-2 RBD-staining of CD19^+^IgD^-^CD27^+^CD38^-^ MBCs at 6 and 12 months in one S-CoV and one M-CoV representative patients. **(B-C)** Frequencies of SARS-CoV-2 RBD-specific cells in live CD19^+^IgD^-^CD27^+^CD38^int/-^ MBCs at 6 (M6) and 12 months (M12) post symptom onset in SARS-CoV-2 recovered patients (S-CoV; dark blue, n=14/14; M-CoV: light blue, n=11/12) (B) and at pre-boost, boost + 7 days and boost + 2 months time points in S-CoV (dark blue, n=17/6/6), M-CoV (light-blue, n=14/21/20) and naive (white, n=10/13/23) patients. Bars indicate mean±SEM. **(D)** UMAP projections of concatenated CD19^+^IgD^-^ B cells multi-parametric FACS analysis from 5 S-CoV and 8 M-CoV patients analyzed longitudinally. His-tagged labeled SARS-CoV-2 RBD-specific cells are overlaid as red dots. **(E-F)** Unsupervised clustering (FlowSOM) performed on the concatenated FACS dataset. Main defined clusters (>2% of total CD19^+^IgD^-^ B cells) are shown as overlaid contour plots on the global UMAP representation (E). Cluster distribution of SARS-CoV-2 RBD-specific cells in identified clusters, at indicated time point, is further displayed as bar plots (F). Bars indicate mean±SEM. **(G)** Representative dot plots for CD71 and CD19 expression in IgD^-^CD19^+^CD38^int/-^ B cells at indicated time points from representative S-CoV, M-CoV and naive patients. SARS-CoV-2 RBD-specific MBCs are overlaid as red dots. **(H)** Frequencies of SARS-CoV-2 RBD-specific cells displaying an activated B cell (CD19^+^CD27^+^IgD^-^CD71^+^) phenotype at indicated time points. **(I)** Proportion of Spike-specific memory B cells recognizing RBD in each individual at the two months time point. Bars indicate mean±SEM. (B) Two-way ANOVA with multiple comparisons of all groups; (C and H) Repeated measures mixed effects model analysis with two sets of multiple comparisons (between donor groups inside each time-point (black lines) and between time points for each donor group (colored lines)) and (I) ordinary one-way ANOVA were performed (Benjamini, Krieger and Yekutieli FDR correction was used for all multiple comparisons). Only significant comparisons are highlighted in panels (C, H, I). (***P<0.0001, *** P<0.001, **P < 0.01, *P < 0.05). See also Figure S2 and Table S2.

To further analyze the impact of mRNA vaccine on the memory B cell compartment in SARS-CoV-2 recovered patients, we performed multi-parametric FACS analysis using the same flow panel which we previously used to describe the initial response against SARS-CoV-2 in these patients (Sokal et al., 2021) and which includes 7 surface markers (CD19, CD21, CD27, CD38, CD71, CD11c, IgD) and His-tagged RBD staining to characterize the main B cell subsets. Unsupervised analysis of CD19^+^IgD^-^ switched B cell populations showed that RBD-specific cells mostly resided in the CD21^+^CD27^+^IgD^-^CD38^int/-^CD71^int/-^ resting memory B cell compartment before vaccination **(Figure 2D-F)**. These cells rapidly switched to a CD27^+^CD38^int/+^CD71^+^ activated B cell phenotype (ABC cluster), in the first 7 days after the boost, together with the emergence of a population of RBD^+^ CD38^high^CD27^high^ ASCs. These activated subsets progressively contracted and matured as resting memory B cells at the latest time point in our study. Atypical memory B cells (DN2; IgD^-^CD27^+/-^CD11c^+^) and ASC precursors (DN1; IgD^-^CD27^-^) RBD-specific clusters were also observed, with a small DN2 fraction persisting up to two months post boost, notably in S-CoV patients **(Figure 2F)**.

The low numbers of RBD-specific B cells in naive patients precluded a robust unsupervised analysis, and we therefore characterized the phenotype of RBD-specific memory B cells from the whole cohort using manual gating, based on the CD19 and CD71 expression profiles, to delineate activated B cells (CD19^high^CD71^+^) and resting MBCs (CD19^+^CD71^low^) among RBD-specific cells over time. A large fraction of RBD-specific MBCs acquired a CD19^high^CD71^+^ activated B cell phenotype in the days following the boost in naive individuals, similarly to M-CoV and S-CoV patients (**Figure 2G-H**), demonstrating that vaccine induced a robust expansion of an RBD-specific activated B cell population in all groups of donors.

The proportion of RBD-specific activated B cells decreased over time, favoring an increase of RBD-specific resting MBCs consistent with the contraction of the vaccine response. This contraction appeared more pronounced in SARS-CoV-2 recovered patients, suggesting the persistence of a GC output in naive individuals (**Figure 2H**). Similar kinetics were observed for S-specific B cells **(Figure S2E-F)**, although it should be noted that RBD-specific B cells still represented a smaller proportion of S-specific cells in naive than in S-CoV individuals at the two-month time point (**Figure 2I**).

### mRNA vaccination maintains the overall diversity of the memory B cell repertoire

To address how mRNA vaccine boost affects the MBC repertoire, we then performed single cell sorting and culture of RBD-specific memory B cells before, 7 days and 2 months after the boost in SARS-CoV-2 recovered patients, and 2 months after the boost in naive individuals **(Figure 3A)**. We obtained a total of 2452 V_H_ sequences from 4 S-CoV, 4-M-CoV and 3 naive individuals (**Table S1 and S2**). Clonally expanded RBD-specific MBC were found before vaccination in all SARS-CoV-2 recovered patients, representing 13-34% of total V_H_ sequences for each donor (**Figure 3B and Figure S3A**). Vaccine-activated MBC showed no evidence of further clonal dominance and conserved their overall diversity, despite the major increase in MBC numbers. The fraction of the repertoire belonging to these clones varies from 19-39%, and remains comparable during contraction, with clones persisting over the pre/post vaccination period observed in all SARS-CoV-2 recovered patients. This overall clonal stability was reflected by similar Shannon entropy values at all studied time points **(Figure S3B)**. Similar clonal expansion was observed in naive individuals (**Figure 3C**), which also harbored high level of convergent RBD-specific clones with SARS-CoV-2 recovered patients based on V_H_ sequences (**Figure 3D**).

**Figure 3.**
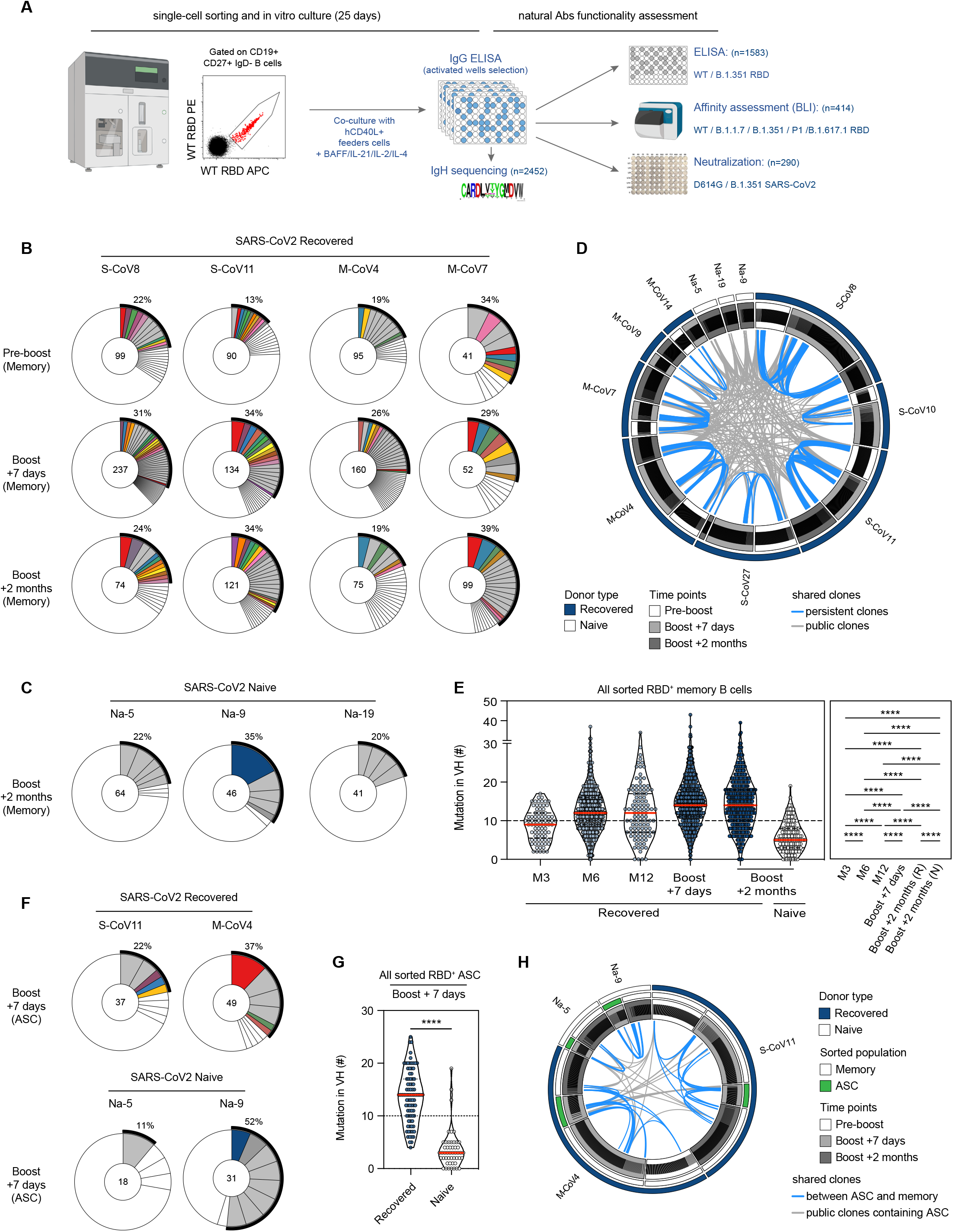
Clonal dynamics of the RBD-specific memory B cell repertoire post-mRNA vaccine boost. **(A)** Experimental scheme for the functional assessment of naturally expressed monoclonal antibodies from RBD-specific memory B cell. **(B-C)** Pie charts representing the clonal distribution of RBD-specific MBCs sorted from 2 S-CoV and 2 M-CoV at pre-boost, boost + 7 days and boost + 2 months time points **(B)** and 2 naive patients at boost + 2 months **(C)**. Clonal representation is depicted according to Wang et al. (2021b) colored slices indicate an expanded MBC clone (2 or more sequences at a given time-point) found at several time-points in the same patient (persistent clone) or in clonal relationship with ASC at boost + 7 days, grey slices indicate an expanded MBC clone found at a single time-point and white slices indicate persistent unique sequences. Outer black semi-circular line indicates the proportion of sequences belonging to expanded clones at a given time point. The total number of sequences is indicated at the pie center. **(D)** Circos plot showing clonal relationships between all sequenced RBD-specific MBCs grouped by donors and time-points. Blue lines connect persistent clones and grey lines connect clones shared by at least two donors (public clones). **(E)** Violin plots showing the number of mutations in the V_H_ segment of RBD-specific MBCs at successive time points in SARS-CoV-2 recovered (M3 = 81 sequences; M6 = 600, M12 = 109; Boost + 7 days = 930, Boost + 2 months = 430) and naive donors (Boost+ 2 months = 151). Red line indicates median. **(F)** Pie chart representing the clonal distribution of RBD-specific ASC sorted for 2 SARS-CoV-2 recovered and 2 naive patients at boost + 7 days. Colored slices indicate a clone in clonal relationship with an expanded MBCs clone from the same donor, grey slices indicate an expanded ASC clone not found in MBCs from the same donor and white slices indicate unique ASC sequences in clonal relationship with a non-expanded MBC clones. Outer black semi-circular line indicates the proportion of sequences belonging to expanded ASC clones. The total number of sequences is indicated at the pie center. (**G)** Violin plots showing the number of mutations in the V_H_ segment of RBD-specific ASCs sorted from 2 SARS-CoV-2 recovered (n= 86 sequences) and 2 naive donors (n= 49 sequences) at boost + 7 days. Red line indicates median. **(H)** Circos plot showing clonal relationships between RBD-specific MBCs (white mid-semi-circular slice) and RBD specific antibody secreting cells (green mid-semi-circular slice) sorted at boost + 7 days from 2 SARS-CoV-2 recovered (one S-CoV, one M-CoV, dark blue outer circular slice) and 2 naive (white outer circular slice). Blue lines indicate shared clones between ASCs and MBCs and grey lines indicate public clones containing ASC. (E-G) Red lines indicate median. Ordinary one-way ANOVA with multiple comparisons (Benjamini, Krieger and Yekutieli FDR correction) (G) and a two-tailed Mann-Whitney test (E) were performed (****P<0.0001). See also Figure S3 and Table S2.

We previously reported, up to 6 months after infection in this cohort of SARS-CoV-2 recovered patients, a progressive accumulation of somatic mutations in RBD-specific clones (Sokal et al., 2021). The mutation level of RBD-specific V_H_ sequences remained stable between 6 and 12 months. In contrast, V_H_ mutation numbers were significantly increased shortly after the boost and this level was maintained in the MBC pool 2 months after (**Figure 3E**). This evolution in mutation profile could further be confirmed at the individual level for 2 patients with complete follow-up from 6 months post symptom onset to 2 months after boost (**Figure S3C-D**). This rapid increase in overall mutational load suggests that a fraction of matured, pre-existing MBCs was selectively mobilized upon vaccine response and persisted over time.

RBD-specific MBCs from naive individuals obtained 2 months after the boost harbored low mutation numbers in their V_H_ genes, consistent with the recruitment of naive B cells in the early phase of the vaccine response. It should moreover be noted that MBCs from naive individuals harbor even lower mutational load than MBCs from SARS-CoV-2 recovered patients at a similar 3 months time point after natural infection (**Figure 3E**). This suggests that natural infection might drive a faster maturation process of the MBC response.

RBD-specific antibody secreting cells (ASC) from SARS-CoV-2 recovered patients sorted 7 days after the boost showed clonal relationships with MBCs and highly mutated V_H_ sequences (**Figure 3F-H and Figure S2A**). In contrast, RBD-specific ASC from naive individuals displayed near germline V_H_ sequences corresponding to the recruitment of naive B cells in the extra-follicular response. Overall, these results show that the activation of the previously matured pool of MBCs by the mRNA vaccine does not bias the clonal diversity of their repertoire in previously infected individuals, while selectively amplifying MBCs with a higher mutation load.

### MBCs clones mobilized by mRNA vaccine contain clones with high-binding affinity against variants of concern

To determine the potency of the MBC compartment mobilized or elicited by mRNA vaccine against the main natural RBD variants, we first performed ELISA assays on 1583 supernatants from single-cell culture of sorted RBD-specific memory B cells from 4 S-CoV, 4-M-CoV and 3 naive individuals **(Figure 4A)**. To normalize for IgG concentration in culture supernatants, we plotted ELISA OD values for WT (Wuhan) vs. B.1.1.7 or B.1.351 RBD variants. Almost all supernatants recognized similarly the WT and B1.1.7 RBD but 20 % of them showed a 10-fold decrease in B.1351 RBD recognition, with similar frequencies of affected supernatants in SARS-CoV-2 recovered and naive individuals (**Figure 4A-B**).

**Figure 4.**
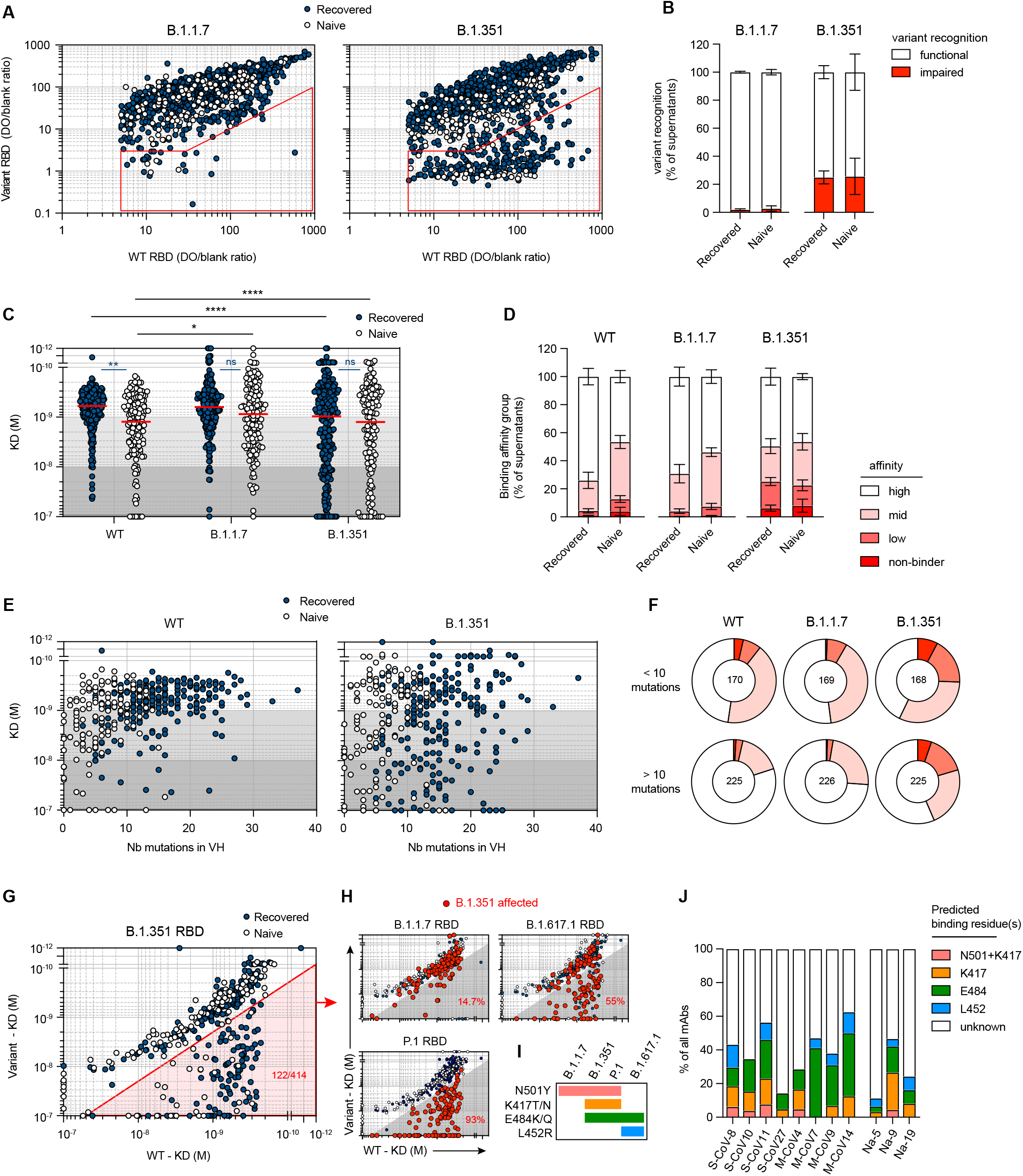
Variant RBD recognition and binding affinity of memory B cells mobilized by the mRNA vaccine boost. **(A**) WT RBD versus B.1.1.7 RBD (left) or B.1.351 RBD ELISA values (right) for all single-cell culture supernatants of RBD-specific memory B cells isolated from SARS-CoV-2 recovered (dark blue, n=952) and naive (white, n=373) donors. Only supernatants with WT ELISA blank-ratio ≥5 are displayed. The red sector identifies naturally expressed antibodies defined as impaired in the recognition of a given variant (variant ELISA blank-ratio <3 or ≥10 fold decrease in variant recognition). **(B)** Frequencies of single RBD-specific memory B culture supernatants with functional or impaired recognition of B.1.1.7 or B.1.351 RBD variants as assessed by ELISA. **(C)** KD (M) measured by bio-layer interferometry for 414 naturally expressed monoclonal antibodies against WT, B.1.1.7 and B.1.351 RBD. Tested monoclonal antibodies were randomly selected from single-cell culture supernatants of RBD-specific MBCs isolated from SARS-CoV-2 recovered (n=267) and naive donors (n=147) and displaying WT RBD ELISA blank ratio ≥3. Background colors define high (KD <10^−9^ M), mid (10^−9^ ≤ KD <10^−8^ M) and low (10^−8^≤ KD <10^−7^) affinity monoclonal antibodies. All monoclonal antibodies with no measurable affinity (KD ≥10^−7^) were considered non-binders. **(D)** Histogram showing the intra-donor binding affinity distribution of monoclonal antibodies tested against WT, B.1.1.7 and B.1.351 RBD variants, as defined in **(C)**, for SARS-CoV-2 recovered or naive donors. Bars indicate mean±SEM. **(E)** Measured KD (M) against WT (left) or B.1.351 RBD (right) versus the number of V_H_ mutations for all tested monoclonal antibodies with available V_H_ sequence from SARS-CoV-2 recovered (dark blue, n=241) and naive (white, n=131)) donors (Spearman correlations for all sequences: V_H_ mutation/WT KD: r = 0.3791, P<0.0001; V_H_ mutation/B.1.351 KD: r = 0.152, P=0.0033). **(F)** Pie chart showing the binding affinity distribution of all tested monoclonal antibodies with low (<10 mutations, upper panel) or high V_H_ mutation numbers (>10, lower panel) against WT, B.1.1.7 and B.1.351 RBD variants as defined in **(C)**. Numbers at center of the pie chart indicate the total number of tested monoclonal antibodies in each group. **(G)** Dot plot representing the KDs for B.1.351 RBD versus WT RBD for all tested monoclonal antibodies from SARS-CoV-2 recovered (dark blue dots) and naive donors (white dots). The red shaded zone indicates B.1.351-affected monoclonal antibodies, defined as those with at least two-fold increased KD for B.1.351 as compared to WT RBD. **(H)** Dot plots representing the KDs for B.1.1.7, B.1.617.1 and P.1 RBD versus WT RBD. B.1.351 affected monoclonal antibodies are highlighted as larger size red dots. Percentages indicate the proportion of B.1.351 affected monoclonal antibodies also affected by each other RBD variants. **(I)** Distribution of known mutations in the RBD domain between B.1.351, B.1.1.7, B.1.617.1 and P.1 SARS-CoV-2 variants. **(J)** Frequencies of predicted essential binding residues, as defined by RBD variants recognition profile in BLI, for all monoclonal antibodies isolated from each of the 11 tested donors. Numbers of tested monoclonal antibodies for each donor are indicated on top of each histogram. (C) A two-way ANOVA with two sets of multiple comparisons (between tested variants inside each group (black lines) and between groups for each tested variants (colored lines)) was performed (Benjamini, Krieger and Yekutieli FDR correction). (****P<0.0001, **P < 0.01, *P < 0.05). See also Figure S4 and Table S2.

We next assayed the affinity of MBCs against B1.1.7, B.1351, P1, and B.1.617 RBD variants using Biolayer interferometry (BLI) in which naturally expressed monoclonal IgG from single-cell culture supernatants were loaded on anti-human IgG biosensors. Monoclonal antibodies expressed by memory B cells from recovered patients were highly enriched for high affinity binders against the WT RBD as compared with naive patients (73.9±5.8% vs 46.5±4.3%, P<0.01) (**Figure 4C-D)**. However, overall affinity profiles were no longer different between recovered and naïve donors when the B.1.351, P1, or B.1.617.1 variants were tested, with approximately 20% of clones with no or low affinity (KD>10^−8^ M) in both groups (**Figure 4C-D and Figure S4A-B**). Somatic hypermutation of V_H_ genes and antigen-driven affinity maturation against the WT RBD accounts for the higher frequency of high-affinity binders (KD<10^−9^ M) in SARS-CoV-2 recovered compared to naive individuals (**Figure 4E-F**). As expected, the number of V_H_ mutations showed reduced correlation with affinity against the B.1.351 RBD (**Figure 4E-F**), in line with the selection of these clones against the WT, and not the mutated RBD, during the GC maturation process.

Two by two comparisons of binding affinities between the WT and the different RBD variants allowed us to validate the sensitivity of our approach with nearly all B.1.351 affected monoclonal antibodies also affected in their binding of the P.1 RBD variant sharing similar set of mutations (namely N501Y, E484K and K417T/N) but not of the B.1.1.7 RBD variant which only harbors the N501Y mutation (**Figure 4H**). Comparison with the more recent B.671.1 RBD variant, which harbor the E484Q but not the K417N/T mutation, further revealed cases of antibodies solely affected in their binding of the B.1.351 and P.1 RBD variants and thus identified as binding the K417 residue (**Figure 4H and Figure S4C-D**). Overall, the natural distribution of mutations in the various VOC RBD (**Figure 4I**) allowed us to predict the identity of key binding amino acid residues within the RBD for 142 out of 414 tested memory B cell-derived monoclonal antibodies **(Figure 4G-H and Figure S4C-D)**. In agreement with recent structural studies (Yuan et al., 2021), clones recognizing the K417 residue were highly enriched for IGHV3-53 and 3-66 genes, whereas clones recognizing the E484K/Q residue were mostly enriched for the IGHV1-2 and 1-69 genes (**Figure S4E**). These results also allowed us to ascribe RBD binding residues of B cells within individual memory repertoires of naive and recovered patients, confirming the major targeting of the E484 residue within the vaccine activated pool (**Figure 4J**), but also highlighting significant inter-donor variability in the overall recognition profile of the SARS-CoV-2 RBD by their memory B cell repertoire.

### The MBC repertoire selected against WT RBD protein upon infection or vaccination contains high-affinity potent B.1.351-neutralizer clones

Finally, to evaluate whether RBD-specific clones mobilized after vaccination, including high affinity B.1.351 RBD binders, display a functional neutralizing activity against the B.1.351 SARS-CoV-2 variant, 279 randomly selected single memory B cell culture supernatants from S-CoV, M-CoV and naive vaccinated individuals were tested in our focus reduction neutralization assay against the D614G and B1.351 SARS-CoV-2 viruses. Potent neutralizing capacity was defined as > 80% of neutralization at 16 nM and weak neutralizing capacity as > 80% of neutralization at 80 nM, with lower values qualified as non-neutralizing. The majority of RBD-specific MBCs clones efficiently neutralized D614G but higher frequency of non-neutralizing clones was observed in naive compared to recovered patients **(Figure 5A and Figure S5A)**. Among potent D614G neutralizers, 40-60% of monoclonal antibodies efficiently neutralized B.1.351, without quantitative difference between M-CoV and naive individuals (**Figure 5B**) or clear association with the V_H_ mutational load or binding affinity to the WT RBD (**Figure 5C and Figure S5A)**. Nonetheless, potent neutralizing memory B cell-derived monoclonal antibodies could be detected in all studied donors, including SARS-CoV-2 naive vaccinees, and represented 20 to 40% of the overall repertoire in most donors.

**Figure 5.**
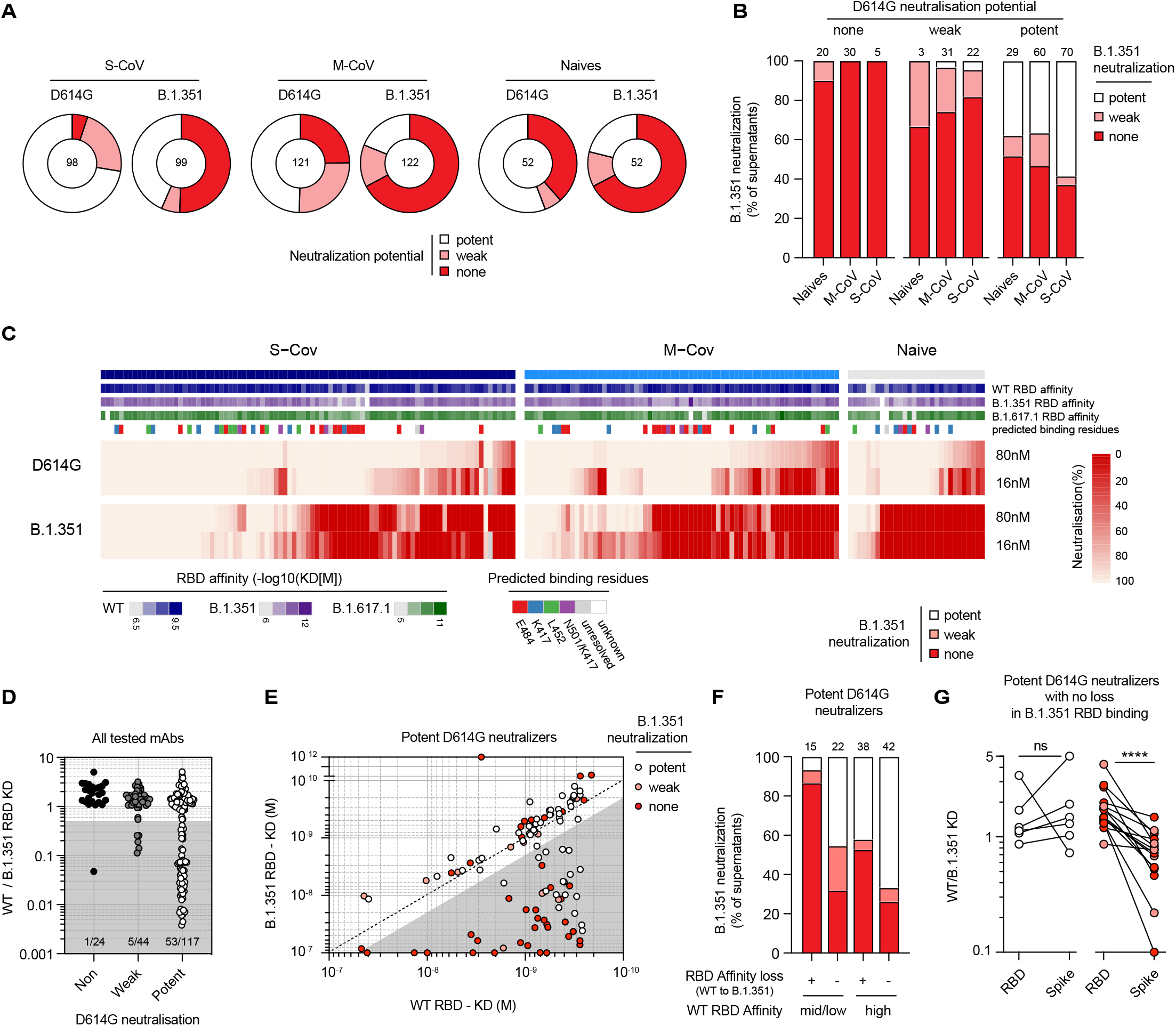
D614G and B.1.351 SARS-CoV-2 neutralization potency of individual memory B cells mobilized by the mRNA vaccine boost. **(A)** Pie charts showing the proportion of single-cell culture supernatants of RBD-specific MBCs isolated from SARS-CoV-2 recovered (S-CoV, n=104; M-CoV, n=123) and naive donors (n=52) displaying potent, weak or no neutralization potential (none) against D614G SARS-CoV-2 and B.1.351 SARS-CoV-2 variants. Potent neutralizers are defined as >80% neutralization at 16 nM, weak neutralizer as neutralization <80% at 16 nM but >80% at 80 nM. Others were defined as non-neutralizing. **(B)** Histogram showing the proportion of B.1.351 SARS-CoV-2 potent, weak and non-neutralizing single-RBD-specific memory B cell culture supernatants grouped based on their neutralization potency against D614G SARS-CoV-2. **(C)** Heatmap showing the in vitro neutralization of D614G SARS-CoV-2 and B.1.351 SARS-CoV-2 at 80 nM and 16 nM for all culture supernatants whose monoclonal antibodies were also tested in BLI (S-CoV, n=91; M-CoV, n=69 and naive, n=30). KD (M) against WT, B.1.351 and B.1.617.1 RBD for each monoclonal antibody are represented on top along with predicted binding residues. **(D)** Ratio of WT RBD KD over B.1.351 RBD KD for all monoclonal antibodies displayed in (C), grouped based on their neutralization potency against D614G SARS-CoV-2 **(E)** KD (M) against B.1.351 versus WT RBD for all D614G SARS-CoV-2 potent neutralizers monoclonal antibodies. Dot color indicates the neutralization potency against B.1.351 SARS-CoV-2 variant. Grey shade highlights binding-impaired clones against the B.1.351 RBD variant as defined in Figure 4G. **(F)** B.1.351 SARS-CoV-2 neutralization potency distribution of all tested potent D614G SARS-CoV-2 neutralizers, grouped based on their affinity for WT RBD and affinity loss against B.1.351. **(G)** WT versus B.1.351 variant RBD or Spike KD ratio for selected monoclonal antibodies showing no clear B.1.351 RBD binding impairment and no loss (left) or clear loss (right) of neutralization potency against the B.1.351 SARS-CoV-2 variant. (G) A paired Wilcoxon test was performed (****P<0.0001). See also Figure S5 and Table S2.

Loss of affinity against B.1.351 RBD mainly affected potent D614G neutralizers **(Figure 5D)**, highlighting the strong evolutionary pressure at the virus level to selectively evade this part of the immune response. Plotting KD values for WT vs. B.1.351 RBD among potent D614G neutralizers further revealed four different profiles correlated with B.1.351 neutralization **(Figure 5E)**. Memory B cell-derived monoclonal antibodies displaying similar affinities against both WT and B.1.351 RBD were enriched in strong neutralizers against the B.1.351 variant **(Figure 5F)** but a subset of these antibodies did display reduced neutralization potency. Interestingly, these monoclonal antibodies showed a selectively reduced affinity against the whole B.1.351 S ectodomain **(Figure 5G)**, suggesting alteration in their binding at the level of the trimeric protein. Among monoclonal antibodies affected in their binding to the B.1.351 RBD (grey zone in **Figure 5E**), maintenance of neutralization activity against the VOC was strongly dictated by original affinity against the WT RBD. Whereas clones with weak affinity against WT RBD were mostly impaired, 40% of clones with high initial affinity against WT RBD remained potent neutralizers against the VOC (white dots in the grey sector, **Figure 5E**). This included monoclonals directed against both the E484 and K417 residues of SARS-CoV-2 RBD (**Figure S5B**). This highlights the key role of affinity maturation in shaping the humoral response and anticipating viral escape.

Overall, these results demonstrate that the MBC pool selected against the WT RBD after mRNA vaccination contains, among its diverse and affinity matured repertoire, a substantial fraction of potent neutralizers against VOC.

## Discussion

Memory B cells display a diverse repertoire allowing for an adaptive response upon re-exposure to the pathogen, especially in the case of variants (Purtha et al., 2011; Weisel et al., 2016). However, repeated antigenic stimulation, either with vaccinations or viral challenge may be deleterious, reducing the diversity of the overall response in which drifted epitopes are less well targeted (Andrews et al., 2015; Mesin et al., 2020). Thus, understanding how mRNA vaccination impacts the memory B cell pool shaped by a previous exposure to SARS-CoV-2 and to determine its capacity to neutralize variants is critical. More generally, to decipher how memory B cells from naive vaccinees differ and evolve in comparison with SARS-CoV-2 recovered patients is also of major importance in the pandemic context.

We report here a longitudinal study of SARS-CoV-2 recovered patients, followed over one year after their initial infection and vaccinated with the BNT162b2 mRNA vaccine. The vaccine response of SARS-CoV-2 naive individuals was analyzed in parallel. The strength of our approach is the large scale, unbiased study of the memory B cell response against SARS-CoV-2 at the single cell level using in vitro activation of randomly sampled memory/activated B cells. This allowed a deep functional analysis which included affinity assessment against 5 RBD variants and determination of neutralization potency of these secreted IgG against two SARS-CoV-2 viruses, without cloning and re-expression intermediates. We focused on the RBD because anti-RBD antibodies contribute to the majority of neutralizing antibodies and its mutations allow for immune escape of VOC (Ju et al., 2020; Robbiani et al., 2020), together with additional targets in the N-terminal domain of the Spike (McCallum et al., 2021).

Our study demonstrates a remarkable stability of the overall RBD-specific memory B cell population up to 12 months after infection with a stable mutation profile, extending observations on memory persistence in COVID-19 (Wang et al., 2021b). Dynamics of RBD-specific cells after mRNA vaccination in SARS-CoV-2 recovered patients reflect the plasticity of the MBC pool, which promptly and widely activates, proliferates and generates ASCs, and then contracts as resting memory B cells (Goel et al., 2021; Reynolds et al., 2021; Wang et al., 2021b). This mobilized population, with a higher mutational load than before vaccinal boost, contains highly-mutated affinity-matured clones that settle, expand and persist for up to 2 months after the boost at a higher level than before. Despite this, longitudinal VDJ sequencing revealed a limited impact on the diversity of this previously matured repertoire. Thus, the expansion of the MBC pool does not seem to impair archives of the B cell specificities that have been selected during the amplification. In contrast, and mirroring early stages of the extrafollicular response after infection (Woodruff et al., 2020), RBD-specific B cells expressing near germline recurrent V_H_ genes with potent affinity are recruited after mRNA vaccination in naive individuals. MBC from naive individuals also acquired somatic mutations with time but harbored, 2 months after the boost, fewer mutations than SARS-CoV-2 recovered patients 3 months after COVID-19. Thus, whereas the MBC pool progressively matures in naive vaccinees, its maturation and amplification were less pronounced, resulting in less RBD-specific MBCs compared to SARS-CoV-2 recovered patients at 12 months after infection (respectively 7.8 and 3 times less than S-CoV and M-CoV) and 2 months after the boost (respectively 18.2 and 6 times less than S-CoV and M-CoV).

As previously reported, most sera of SARS-CoV-2 recovered patients efficiently neutralized the B1.351 variant after mRNA vaccine, which contrasted with the significant lower neutralization potency of the sera of naïve vaccinated individuals (Goel et al., 2021; Reynolds et al., 2021; Stamatatos et al., 2021; Wang et al., 2021b). This likely reflects the quality and maturation of the B cell pool mobilized by the vaccine, i.e. an extrafollicular response in naive vaccinees and a mature MBC response in recovered patients.

A fraction of recurrent and convergent V_H_ genes of memory B cells failed to recognize the B.1351 variant RBD in both cohorts but a large proportion of clones retained high affinity against this variant. This is consistent with a recent report showing that B cell clones expressing potent antibodies are selectively retained in the repertoire over time (Wang et al., 2021b). This phenomenon occurs independently of the mutational load, high-affinity clones against variants being “randomly” present in the memory repertoire of SARS-CoV-2 recovered or naive individuals. So, despite the fact that amplitude and quality of the MBC response after mRNA vaccine appears to be lower in naive than in previously infected individuals, high-affinity clones with neutralizing potency against VOC settle in their repertoire, suggesting that their MBC pool could compensate for the time-dependent decay of the initial antibody response.

Correlation of neutralization and affinity shows a complex profile for antibodies expressed by memory B cells. Immune escape can affect high-affinity clones against WT RBD, but at the same time, a proportion of clones with high affinity for WT RBD maintain neutralizing potency against B1.351, contrasting with clones with low or weak affinity that constantly failed to neutralize variants. It indicates that high affinity provides some flexibility for antibodies to cope with mutations affecting their binding target and to conserve their neutralization potency. Consistent with structural analysis (Yuan et al., 2021), determination of targeted epitopes using binding of different RBD of VOC shows that E484 preferentially affected the binding affinity of the IGHV1-2/1-69 genes, and K417N and N501Y the one of IGHV3-53/3-66 and IGHV1-2. It underlines, that if particular antibody lineages are affected by RBD mutations, others may retain neutralizing properties (Barnes et al., 2020; Greaney et al., 2021b, 2021c; Muecksch et al., 2021; Scheid et al., 2021; Wang et al., 2021b).

Altogether, these data describe an immune response maturing with time in SARS-CoV-2 convalescent patients, and resulting in a massive, high-affinity response after vaccination, which, even imprinted by the Wuhan-type RBD, displays an improved recognition of the RBD variants as well. In this immune evolution scheme, the response of naive vaccinees nevertheless lags behind the maturation process that took place during infection. Our observations suggest that repeated challenges, even against the original spike protein, will compensate for these differences by recall of affinity-matured memory B cells and allow vaccinated people to cope efficiently with most variants actually described.

## Acknowledgments

We thank Garnett Kelsoe for providing us with the human cell culture system, together with invaluable advices. We thank Fabrice Agou and Alix Boucharlat (Chemogenomic and Biological Screening Platform, Center for Technological Resources and Research (C2RT), Institut Pasteur, Paris, France) for advice on the use of the Octet HTX instrument for BLI measurements. We also thank Sébastien Storck, Lucie Da Silva, Sandra Weller for their advices and support; we thank the physicians, Constance Guillaud, Raphael Lepeule, Frédéric Schlemmer, Elena Fois, Henri Guillet, Nicolas De Prost, Pascal Lim, whose patients were included in this study.

## Author contributions

Conceptualization: P.C., A.S., JC.W., CA.R., and M.M.; Data curation: P.C., A.S., M.B., G.BS, I.A. Formal Analysis: A.S., P.C., M.B., G.BS, I.A., M.Bo.; A.DLS, A.V., S.F., I.F.; Funding acquisition: S.F., JC.W., CA.R., and M.M.; Investigation: A.S., P.C., A.DLS., I.A., A.V.; Methodology: A.S., JC.W., CA.R., P.C. and M.M.; Project administration: P.C., F.CP., JC.W, CA.R., and M.M.; Resources: JM.P., F.NP., S.F., E.C., L.G., Ma.Mi., B.G., S.G., G.M., Y.NG., V.Z., P.B., F.R., CA.R., M.M.; Software: P.C; Supervision: P.C., JC.W., CA.R, P.B., F.R. and M.M.; Validation: A.S., I.A., A.V., A.DLS, M.B., CA.R.,PC., MM. ; Visualization: P.C., A.S., I.A., A.DLS., M.M.; Writing – original draft: P.C., A.S., JC.W, CA.R., and M.M.; Writing – review & editing: all authors.

## Methods

| Reagent or resource | Source | Identifier |
| --- | --- | --- |
| **Antibody** |  |  |
| CD3 | Biologend | UCHT1 ; Cat#300425 ; RRID : AB_830754 |
| CD14 | BD Bioscience | M φP9 ; Cat#561709; RRID : AB_1645464 |
| CD19 | BD Bioscience | HIB19 ; Cat# 562321 ; RRID : AB_11154408 |
| CD38 | BD Bioscience | HIT2 ; Cat# 551400 ; RRID AB_394184 |
| CD27 | Biologend | M-T271 ; Cat# 356417 ; RRID: AB_2562598 |
| CD11c | BD Bioscience | S-HCL-3 ; Cat#744436 ; RRID : AB_2742232 |
| IgD | Life technologies | Polyclonal ; Cat# H15501 ; RRID : AB_2536563 |
| His Tag | Biologend | J095G46 ; Cat# 362605 ; RRID:AB_2563634 |
| His Tag | Biologend | J095G46 ; Cat# 362603 ; RRID:AB_2715818 |
| CD71 | Biologend | CY1G4 ; Cat# 334111 ; RRID:AB_2563118 |
| CD21 | BD Bioscience | B-ly4; Cat#563163 ; RRID:AB_2741028 |
| N SARS-CoV-2 | Institut Pasteur | Rabbit polyclonal (N.Escriou) |
| Anti-Human Fc Capture Biosensors | Sartorius | Cat#18-5060 |
| **Biological samples** |  |  |
| Cryopreserved PBMCs from S-Cov, M-Cov and naive patients | Henri Mondor Hospital, Assistance Publique des Hôpitaux de Paris |  |
| D614G SARS-CoV-2 virus (hCoV-19/France/GE1973/2020) | Institut Pasteur | CNR Respiratory Viruses (S.Van der Werf) |
| B.1.351 SARS-CoV-2 virus (CNR 202100078) | Institut Pasteur | CNR Respiratory Viruses (S.Van der Werf) |
| **Chemical, peptides, and recombinant proteins** |  |  |
| SARS-CoV-2 Spike | Institut Pasteur | Virologie Structurale (F. Rey) |
| B.1.351 SARS-CoV-2 Spike | Institut Pasteur | Virologie Structurale (F. Rey) |
| WT SARS-CoV-2 RBD | Institut Pasteur | Virologie Structurale (F. Rey) |
| B.1.1.7 SARS-CoV-2 RBD | Institut Pasteur | Virologie Structurale (F. Rey) |
| B.1.351 SARS-CoV-2 RBD | Institut Pasteur | Virologie Structurale (F. Rey) |
| P.1 SARS-CoV-2 RBD | Institut Pasteur | Virologie Structurale (F. Rey) |
| B.1.617.1 SARS-CoV-2 RBD | Institut Pasteur | Virologie Structurale (F. Rey) |
| Live dead aqua | Life technologies | L34957 |
| Recombinant human IL-2 | PeproTech | 200-02 |
| Recombinant human IL-4 | PeproTech | 200-04 |
| Recombinant human IL-21 | PeproTech | 210-21 |
| Recombinant human BAFF | PeproTech | 310-13 |
| **Softwares and algorithms** |  |  |
| Kaluza v2.1 | Beckman Coulter | https://www.beckman.fr |
| Flowjo v10.7.1 | FlowJo, LLC | https://www.flowjo.com |
| GraphPad Prism v9 | GraphPad | https://www.graphpad.com |
| Codon Code Aligner v9 | Codon Code Corporation | https://www.codoncode.com/ |
| R v4.0.2 | R Foundation | https://www.r-project.org |
| RStudio v1.3.1056 | RStudio | https://rstudio.com |
| IgBLASTn v1.16.0 | NCBI | https://www.ncbi.nlm.nih.gov/igblast/ |
| **HT Data analysis software 11.1** | ForteBio | https://www.sartorius.com |
| Adobe Illustrator (CS6) | Adobe | https://www.adobe.com |

## RESOURCE AVAILABILITY

### Lead Contact

Further information and requests for resources and reagents should be directed to and will be fulfilled by the Lead Contact, Matthieu Mahévas.

### Materials Availability

No unique materials were generated for this study.

### Data and Code Availability

## EXPERIMENTAL MODEL AND SUBJECT DETAILS

### Study participants

In total, 34 patients with recovered COVID-19 from the original MEMO-COV-2 cohort were followed up to 12 months post-infection and/or vaccination. Among them, 17 patients had severe COVID-19 (patients requiring oxygen, S-CoV) and 17 had a mild COVID-19 disease (mainly healthcare workers, M-CoV). An additional cohort of 9 patients who experienced mild COVID-19 during the first wave and were vaccinated at least six months after the infection were recruited. SARS-CoV-2 infection was defined as confirmed reverse transcriptase polymerase chain reaction (RT-PCR) on nasal swab or clinical presentation associated with typical aspect on CT-scan and/or serological evidence. Twenty-five healthcare workers who had no history of COVID-19 and negative IgG anti-nucleocapsid (and/or Spike) were enrolled in the naive group (IRB 2018-A01610-55).

All vaccinated subjects received the BNT162b2 mRNA vaccine. SARS-CoV-2 recovered patients received only one dose, in line with French guidelines, except 3 who received 2 doses (See **Table S1**). First injection was realized in mean 309 days (± SD 44.6 days) after the infection. Naive patients received two doses at a mean 27.7 days (± SD 1.8 days) interval. Prior to vaccination, samples were collected from SARS-CoV-2 recovered patients 12 months post symptoms onset (mean ±SD 329.1 ± 15: 8.8 days after disease onset for S-CoV, and 342.0 ± 8.6 days after disease onset for M-CoV). Samples at 12 months post disease onset were defined as “pre-boost”. For patients not sampled before vaccination (n=9/34), sample at 6 months was considered as “pre-boost”. For naive patients, the “prime” time-point was defined as the sampling between the two doses and was drawn at a mean 20.2 ± 5.9 days after the first injection.

Samples were additionally collected shortly after the boost (mean ±SD: 10 ± 5.3 days for S-CoV; 23 ± 6.1 days for M-CoV and 9 ± 4.0 days for naive), and 2 months after the boost (mean ±SD: 64.7 ± 15.3 days for S-CoV ; 63.2 ± 11.9 days for M-CoV and 63.3 ± 9.0 days for naive). Clinical and biological characteristics of these patients are summarized in **Table S1**. Patients were recruited at the Henri Mondor University Hospital (AP-HP), between March and April 2021. MEMO-COV-2 study (NCT04402892) was approved by the ethical committee Ile-de-France VI (Number: 40-20 HPS), and was performed in accordance with the French law. Written informed consent was obtained from all participants.

## METHOD DETAILS

### Anti-RBD (S) SARS-CoV-2 antibodies assay

Serum samples were analyzed for IgG anti-S-RBD IgG titration with the SARS-CoV-2 IgG Quant II assay (ARCHITECT®, Abbott Laboratories). The latter assay is an automated chemiluminescence microparticle immunoassay (CMIA) that quantifies anti-RBD IgG, with 50 AU/mL as a positive cut-off and a maximal threshold of quantification of 40,000 AU/mL. All assays were performed by trained laboratory technicians according to the manufacturer standard procedures.

### Recombinant protein purification

#### Construct design

The ectodomain from the SARS-CoV-2 Spike (residues 1-1208) was designed as a stabilized construct with six proline mutations (F817P, A892P, A899P, A942P, K986P, V987P), a GSAS substitution at the furin cleavage site (residues 682–685) and a C-terminal Foldon trimerization motif (Hsieh, 2020), followed by Hisx8, Strep and Avi tags. This construct was cloned using its endogenous signal peptide in pcDNA3.1(+).

The SARS-CoV-2 Receptor Binding Domain (RBD) was cloned in pcDNA3.1(+) encompassing the Spike (S) residues 331-528, and it was flanked by an N-terminal IgK signal peptide and a C-terminal Thrombin cleavage site followed by Hisx8, Strep and Avi tags. The mutations present on the B.1.1.7 (N501Y), B.1.351 (K417N, E484K, N501Y), P.1 (K417T, E484K, N501Y) and B.1.617.1 (L452R, E484Q) variants were introduced by PCR mutagenesis using standard methods.

#### Protein expression and purification

The plasmids coding for recombinant proteins were transiently transfected in Expi293F™cells (Thermo Fischer) using FectoPRO® DNA transfection reagent (Polyplus), according to the manufacturer’s instructions. The cells were incubated at 37 °C for 5 days and then the culture was centrifuged and the supernatant was concentrated. The proteins were purified from the supernatant by affinity chromatography using StrepTactin columns (IBA) (SARS-CoV-2 S) or His-Trap™Excel columns (Cytiva) (SARS-CoV-2 RBD). A final step of size-exclusion chromatography (SEC) in PBS was also performed, using either a Superose6 10/300 column (Cytiva) for the SARS-CoV-2 S, or a Superdex200 10/300 (Cytiva) for the SARS-CoV-2 RBD.

### Flow cytometry and cell sorting

PBMCs were isolated from venous blood samples via standard density gradient centrifugation and used after cryopreservation at -150°C. Cells were thawed in RPMI-1640 (Gibco)-10% FBS (Gibco), washed twice and incubated with 5 µg of the SARS-CoV-2 His-tagged spike protein or WT RBD in 100 µL of PBS (Gibco)-2% FBS during 20 min on ice. Cells were washed and resuspended in the same conditions, then the fluorochrome-conjugated antibody cocktail including the 2 anti-His antibodies was added at pre-titrated concentrations for 20 min at 4°C and viable cells were identified using a LIVE/DEAD Fixable Aqua Dead Cell Stain Kit (Thermo Fisher Scientific) incubated with conjugated antibodies. Samples were acquired using a LSR Fortessa SORP (BD Biosciences). For cell sorting, cells were stained using the same protocol and then sorted in 96 plates using the ultra-purity mode on a MA900 cell sorter (SONY), or Aria III (BD Biosciences). Data were analyzed with FlowJo or Kaluza softwares. Detailed gating strategies for individual markers are depicted in Figure S1.

For UMAP generation and visualization (**Figure 2**), FCS files from 5 S-CoV, 8 M-CoV and 4 naive patients with complete panel acquisition at pre boost, boost + 7 days and boost + 2 months (**Table S1**) were concatenated. The UMAP (v3.1) plugin in FlowJO was used to calculate the UMAP coordinates for the resulting 536.161 cells (with 30 neighbors, metric = euclidian and minimum distance = 0.5 as default parameters). The FlowSOM (v2.6) plugin was used in parallel on the same downsampled dataset to create a self-organizing map (using n = 10 clusters as default parameter). This self-organizing map was then applied to the initial FCS files to calculate both total and RBD-specific memory B cell repartition in identified clusters on all collected cells for each donor. Naive patients were not further analyzed due to the very low number of RBD specific B cells. Both UMAP and FlowSOM plugin were run taking into account fluorescent intensities from the following parameters: FSC-A, SSC-A, CD19, CD21, CD11c, CD71, CD38, CD27 and IgD. Contour plots (equal probability contouring, with levels set to 5% of gated populations) for each identified cluster were further overlaid on UMAP projection in FlowJO. For visualization purposes, only the outermost density representing 95% of the total gated cells was kept for the final figure, all other levels were removed in Adobe Illustrator.

### Single-cell culture

Single cell culture was performed as previously described (Crickx et al., 2019): single B cells were sorted in 96-well plates containing MS40 cells expressing CD40L (kind gift from G. Kelsoe). Cells were co-cultured at 37°C with 5% CO2 during 21 or 25 days in RPMI-1640 (Invitrogen) supplemented with 10% HyClone FBS (Thermo Scientific), 55 µM 2-mercaptoethanol, 10 mM HEPES, 1 mM sodium pyruvate, 100 units/mL penicillin, 100 µg/mL streptomycin, and MEM non-essential amino acids (all Invitrogen), with the addition of recombinant human BAFF (10 ng/ml), IL2 (50 ng/ml), IL4 (10 ng/ml), and IL21 (10 ng/ml; all Peprotech). Part of the supernatant was carefully removed at days 4, 8, 12, 15 and 18 and the same amount of fresh medium with cytokines was added to the cultures. After 21 days of single cell culture, supernatants were harvested and stored at -20°C. Cell pellets were placed on ice and gently washed with PBS (Gibco) before being resuspended in 50 µL of RLT buffer (Qiagen) supplemented with 10% 2-mercaptoethanol and subsequently stored at -80°C until further processing.

### ELISA

Total IgG and SARS-CoV-2 WT RBD, B.1.1.7 RBD and B.1.351 RBD-specific IgG from culture supernatants were detected by home-made ELISA. Briefly, 96 well ELISA plates (Thermo Fisher) were coated with either goat anti-human Ig (10 μg/ml, Invitrogen) or recombinant SARS-CoV-2 WT RBD, B.1.1.7 RBD or B.1.351 RBD protein (2.5 µg/ml each) in sodium carbonate during 1h at 37°C. After plate blocking, cell culture supernatants were added for 1hr, then ELISA were developed using HRP-goat anti-human IgG (1 μg/ml, Immunotech) and TMB substrate (Eurobio). OD450 and OD620 were measured and Ab-reactivity was calculated after subtraction of blank wells. Supernatants whose ratio of OD450-OD620 over control wells (consisting of supernatant from wells that contained single-cell sorted spike-negative memory B cells from the same single cell culture assay) was over 3 were considered as positive for WT RBD, B.1.1.7 RBD or B.1.351 RBD ELISA. PBS was used to define background OD450-OD620.

### Single-cell IgH sequencing

Clones whose culture had proven successful (IgG concentration ≥ 1 µg/mL at day 21-25) were selected and extracted using the NucleoSpin96 RNA extraction kit (Macherey-Nagel) according to the manufacturer’s instruction. A reverse transcription step was then performed using the SuperScript IV enzyme (ThermoFisher) in a 14 μl final volume (42°C 10 min, 25°C 10 min, 50°C 60 min, 94°C 5 min) with 4 µl of RNA and random hexamers (Thermofisher scientific). A PCR was further performed based on the protocol established by Tiller et al (Tiller et al., 2008). Briefly, 3.5 μl of cDNA was used as template and amplified in a total volume of 40 μl with a mix of forward L-VH primers (Table S4) and reverse Cγ primer and using the HotStar® Taq DNA polymerase (Qiagen) and 50 cycles of PCR (94°C 30 s, 58°C 30 s, 72°C 60 s). PCR products were sequenced with the reverse primer CHG-D1 and read on ABI PRISM 3130XL genetic analyzer (Applied Biosystems). Sequence quality was verified with the CodonCode Aligner software (CodonCode Corporation).

For specific patients and time points (see **Table S1**), some IgH sequences were obtained directly from single-cell sorting in 4 µL lysis buffer containing PBS (Gibco), DTT (ThermoFisher) and RNAsin (Promega). Reverse transcription and a first PCR was performed as described above (50 cycles) before a second 50-cycles PCR using 5’AgeI VH primer mix and CHG-D1 3’ primer, before sequencing.

### Computational analyses of VDJ sequences

Processed FASTA sequences from cultured single-cell V_H_ sequencing were annotated using Igblast v1.16.0 against the human IMGT reference database. Clonal cluster assignment (DefineClones.py) and germline reconstruction (CreateGermlines.py) was performed using the Immcantation/Change-O toolkit (Gupta et al., 2015) on all heavy chain V sequences. Sequences that had the same V-gene, same J-gene, including ambiguous assignments, and same CDR3 length with maximal length-normalized nucleotide hamming distance of 0.15 were considered as potentially belonging to the same clonal group. Mutation frequencies in V genes were then calculated using the calcObservedMutations() function from the Immcantation/SHazaM v1.0.2 R package. VH repartitions and Shannon entropies were calculated using the countGenes() and alphaDiversity() functions from the Immcantation/alakazam v1.1.0 R package. Further clonal analyses on all productively rearranged sequences were implemented in R. Graphics were obtained using the ggplot2 v3.3.3, pheatmap v1.0.12 and circlize v0.4.12 packages.

### Avidity measurement using biolayer interferometry (Octet)

Biolayer interferometry assays were performed using the Octet HTX instrument (ForteBio). Anti-Human Fc Capture (AHC) biosensors (18-5060) were immersed in supernatants from single-cell memory B cell culture (or control monoclonal antibody) at 25°C for 1800 seconds. Biosensors were equilibrated for 10 minutes in buffer prior to measurement. Association was performed for 600 s in PBS-BT with WT or variant RBD at 100nM followed by dissociation for 600s in PBS-BT. Biosensor regeneration was performed by alternating 30s cycles of regeneration buffer (glycine HCl, 10 mM, pH 2.0) and 30s of PBS-BT for 3 cycles. Traces were reference sensor subtracted and curve fitting was performed using a local 1:1 binding model in the HT Data analysis software 11.1 (ForteBio). Sensors with response values below 0.1nm were excluded or considered non-binding. For binding clones, only those with full R^2^>0.8 were retained for further analysis.

### Virus strains

The reference D614G strain (hCoV-19/France/GE1973/2020) and the B.1.351 strain (CNR 202100078) were supplied by the National Reference Centre for Respiratory Viruses hosted by Institut Pasteur and headed by Sylvie van der Werf. The viral strains were supplied through the European Virus Archive goes Global (EVAg) platform, a project that has received funding from the European Union’s Horizon 2020 research and innovation program under grant agreement number 653316.

D614G and B.1.351 viral stocks were prepared by amplification and titration in Vero E6 cells and were used at passage 3 and passage 2 respectively. Single use aliquots stored at -80°C were used for all the assays.

### Virus neutralization assay

Virus neutralization was evaluated by a focus reduction neutralization test (FRNT). Vero E6 cells were seeded at 2×10^4^ cells/well in a 96-well plate 24h before the assay. Two-hundred focus-forming units (ffu) of each virus were pre-incubated with serial dilutions of heat-inactivated sera or B-cell clone supernatants for 1hr at 37°C before infection of cells for 2hrs. The virus/antibody mix was then removed and foci were left to develop in presence of 1.5% methylcellulose for 2 days. Cells were fixed with 4% formaldehyde and foci were revealed using a rabbit anti-SARS-CoV-2 N antibody (gift of Nicolas Escriou) and anti-rabbit secondary HRP-conjugated secondary antibody. Foci were visualized by diaminobenzidine (DAB) staining and counted using an Immunospot S6 Analyser (Cellular Technology Limited CTL). B-cell culture media and supernatants from RBD negative clones were used as negative control. Pre-pandemic serum (March 2012) was used as negative control for sera titration and was obtained from an anonymous donor through the ICAReB platform (BRIF code n°BB-0033-00062) of Institut Pasteur that collects and manages bioresources following ISO (International Organization for Standardization) 9001 and NF S 96-900 quality standards. Percentage of virus neutralization was calculated as (100 -((#foci sample / #foci control)*100)). Sera IC50 were calculated over 6 four-fold serial dilutions from 1/10 to 1/10000 using the equation log (inhibitor) vs. normalized response – Variable slope in Prism 9 (GraphPad software LLC). For some of the sera (namely the infection/post-vaccination group), we were not able to obtain a complete neutralization curve and set up an arbitrary IC50 cut off of >1/2560 sera dilution. Two IgG concentrations (80 nM and 16 nM) were tested for each sample and each virus.

### Statistics

Ordinary One-way ANOVA, Two-way ANOVA, Repeated measures mixed effects model analysis, Kruskal-Wallis test and Mann-Whitney test were used to compare continuous variables as appropriate (indicated in Figures). Benjamini, Krieger and Yekutieli FDR correction was used for all multiple comparisons. A *P*-value ≤ 0.05 was considered statistically significant. Statistical analyses were all performed using GraphPad Prism 9.0 (La Jolla, CA, USA).

